# GDF8 Contributes to the Adverse Effects of Liver Injury on Skeletal Muscle Homeostasis and Regeneration

**DOI:** 10.1101/2021.04.27.441640

**Authors:** Alexander Culver, Matthew Hamang, Yan Wang, Emily White, Samer Gawrieh, Raj K. Vuppalanchi, Naga P. Chalasani, Guoli Dai, Benjamin C. Yaden

## Abstract

**Background:** An emerging clinical phenomenon in patients with end stage liver disease is progressive skeletal muscle atrophy. This loss in lean mass predicts poor survival outcomes for liver disease patients and highlights an underappreciated crosstalk between injured liver and muscle that lacks defined mediators. The purpose of our study was to identify potential liver-muscle mediator(s) in pre-clinical *in vivo* models of liver injury which may contribute to the muscle loss observed in liver disease.

**Methods:** Utilizing a mouse model of carbon tetrachloride CCl_4_-induced liver injury in the presence or absence of cardiotoxin-induced muscle injury, we evaluated whether neutralizing Activin type IIB receptor (ActRIIB) ligands, or specifically growth differentiation factor 8 (Gdf8), could preserve or reverse muscle atrophy associated with liver disease.

**Results:** We found that hepatic injury via CCl_4_ or bile duct ligation (BDL) similarly caused significant muscle atrophy along with decreased gene expression in key myogenesis markers. This adverse effect of injured liver on muscle were completely prevented and reversed by the intervention of Activin type IIB receptor (ActRIIB)-Fc fusion protein, which neutralizes the ActRIIB ligands, including Activins and growth differentiation factor 8 (Gdf8 or myostatin). The results indicate that ActRIIB ligands promoted muscle atrophy which was manifested in response to hepatic injury/disease and conferred the negative communication of injured liver with muscle. Indeed, direct injection of exogenous Gdf8 protein into muscle along with acute focal muscle injury recapitulated similar dysregulated muscle regeneration as observed with liver injury. Furthermore, we found that hepatocytes produced Gdf8 in response to liver injury in rodents and in patients with end stage liver disease. A neutralizing antibody to Gdf8 attenuated muscle atrophy and unexpectedly ameliorated liver fibrosis in both CCl_4_ and BDL models. Following this observation, we demonstrated Gdf8’s ability to induce fibrogenesis in stellate cells, potentially identifying a novel hepatic role for this protein. Moreover, hepatic Gdf8 promoted muscle wasting in response to liver damage and hindered skeletal muscle regeneration.

**Conclusion:** Our findings identified Gdf8 as a novel hepatomyokine contributing to injured liver-muscle negative crosstalk and liver injury progression. Moreover, we demonstrated a promising therapeutic strategy for muscle atrophy accompanying liver diseases.

## INTRODUCTION

The liver and skeletal muscle systems have a well-established interdependent endocrinological connection in both physiological and pathological states. One example of this inter-dependency is observed through their mutual contribution to the maintenance of glucose homeostasis through storage and metabolic mechanisms. In contrast, patients with liver cirrhosis exhibit abnormal capacity to store glucose as glycogen in skeletal muscle [1]. Moreover, skeletal muscle wasting, or sarcopenia is recognized as a major complication in patients with cirrhosis, non-alcoholic fatty liver disease (NAFLD) and the more severe form known as nonalcoholic steatohepatitis (NASH) [2,3]. Recent clinical studies demonstrate a linear association of sarcopenia with severity of liver fibrosis in NASH patients [4–6]. In addition, the degree of muscle wasting is a strong correlate to adverse clinical outcomes and follow-up hospital costs in these patients [4, 7-12]. Patients with cirrhotic livers and concomitant muscle atrophy tend to have lower survival rates and more post-liver transplant complications. Paradoxically, most complications in liver cirrhosis patients resolve following a successful transplant, except for their underlying muscle wasting [13]. Thus, the reversal of muscle mass loss is paramount for the clinical management of these patients [14–16]. Unfortunately, effective therapies are lacking because the signaling mediators of the liver-muscle axis remain unclear [17]. Therefore, it is a priority to identify and delineate the systemic cues that initiate and promote this sequela of liver diseases.

The TGFβ superfamily of secreted proteins include activins, growth differentiation factors (Gdfs), and bone morphogenetic proteins (BMPs). They engage with Activin type II and type I receptors complexes to initiate Smad signaling to modulating organ development, growth, homeostasis, and repair [17–19]. Notably, it has been proposed that members of the TGFβ superfamily play integral roles in homeostasis and disease states of both liver and skeletal muscle and certain members may even serve as a nexus between the two organs [20–22]. In the present study, we set out to determine (1) whether TGFβ ligands are causal mediators responsible for the deleterious communication between the injured liver with normal or injured skeletal muscle. (2) We also evaluate the potential to prevent or reverse liver injury-induced muscle atrophy by inhibiting TGFβ superfamily signaling. To test this hypothesis, we utilized a combination of fusion proteins (ActRIIB-Fc, ActRIIA-Fc) and neutralizing antibodies to activin A and myostatin (Gdf8) to demonstrate the importance of this signaling axis in this pre-clinical setting.

ActRIIB-Fc is a fusion protein consisting of the extracellular ligand-binding domains of Activin type IIB receptor with the Fc portion of mouse immunoglobulin G (IgG). ActRIIB-Fc primarily binds to and inhibits Activins, Gdf8, and potentially other TGFβ superfamily members [23]. We found that ActRIIB-Fc treatment exhibits profound preventive and therapeutic effects on muscle atrophy and regeneration defects concomitant to liver injury largely by neutralizing Gdf8.

## METHODS

### Approval of animal studies

All mouse experiments are performed under the approval of Eli Lilly’s Institutional Animal Care and Use Committee and are in accordance with the NIH guide for the care and use of laboratory animals. For all studies, ten to twelve-week-old C57BL/6 or ICR mice were used (Envigo, Indianapolis, IN).

### Liver fibrosis models

Hepatic fibrosis was induced by 10 mL/kg intraperitoneal injections of 1:10 diluted carbon tetrachloride (CCl_4_) (Sigma Aldrich, St. Louis, MO) in corn oil twice weekly for a total of 6 to 9 weeks for chronic studies. For acute studies, tissues were harvested 6, 24, or 48 hours following a single injection of CCl_4_. Bile duct ligation was carried out as previously described [24, 25]. Under isoflurane anesthesia (2 to 4 vol %) male ICR mice were placed on a heat pad and laparatomized. The common bile duct was exposed, isolated, ligated two times with non-resorbable sutures (Catgut, Markneukirchen, Germany). Sham-operated mice underwent a laparotomy with exposure but not ligation of the bile duct. The abdominal muscle and skin layers were stitched, and the mice were treated with ketoprofen as an analgesic.

### Skeletal muscle injury model

Muscle injury was induced as previously described [26], with slight modifications. Briefly, muscle injury was induced by a 100 μL injection of a 10 μM cardiotoxin (CTX) (Sigma-Aldrich, St. Louis, MO; part #C3987) solution into the gastrocnemius muscle with a three point injection technique to fully cover the lateral and medial gastrocnemius.

### Gene expression analysis

RNA was extracted from isolated tissues using TRIzol reagent (Life Technologies, Grand Island, NY). Total RNAs were reverse transcribed using the High Capacity cDNA Archive Kit (Applied Biosystems, Foster City, CA). All cDNAs were assayed for genes of interest using TaqMan Gene Expression Analysis (Applied Biosystems) and quantified by 2-ΔΔCt method. In situ hybridization was performed on FFPE tissue sections using mouse and human Gdf8 RNAscope probes according to the manufacturer’s protocol (Advanced Cell Diagnostics).

### Protein quantification

Tissue lysates were generated using 1 mL per 100 mg tissue in lysis buffer (Cell signaling Technologies, Danvers, MA). Total protein contents were quantified using a Pierce BCA Protein Assay Kit (Thermo Scientific). Gdf8 and Smad2/3 proteins were quantified by ELISA methods (GDF8/Myostatin Quantikine ELISA kit, Cat # DGDF80, R&D Systems, Minneapolis, MN; PathScan phospho-Smad2 Ser465/467/Smad3 ser423/425 and total for Smad 2 and 3, kits #12001, 1200, 12002, 7244, Cell Signaling, Danvers, MA). Procedures were followed according to the manufacturer’s guidelines.

### Histology and immunohistochemistry

Muscle or liver tissue was evaluated using hematoxylin and eosin (H&E) or Masson’s trichrome staining (Abcam, Cambridge, MA). For each muscle, distribution of the fiber diameter was calculated by analyzing ~200 myofibers using digital slide scanning (ScanScope XT, Aperio, Vista, CA). Collagen proportionate area (CPA) was quantified using the HALO Image Analysis Platform (Albuquerque, NM) of liver sections stained with Masson’s trichrome. FFPE liver sections were assayed for Gdf8 content via a commercial antibody (LSBio, LS-C37420/170820).

### Human liver sample collection

Liver tissue was collected from individuals with established cirrhosis awaiting liver transplantation at the time of their liver transplantation procedure in the operating room. Demographic data, etiology of cirrhosis, and other relevant information such as medication, alcohol use, and smoking history were captured at the time of enrollment. Liver tissue samples were snap frozen using liquid nitrogen and stored at −80° C until use. All samples were collected and handled equally except for duration of storage. This study was reviewed and approved by the Institutional Review Board (IUPUI IRB: EX0904-11).

### Data analysis and presentation

All results were expressed as mean ± standard errors mean (SEM). Significance (p-value ≤ 0.05) of various parameters was analyzed by one-way ANOVA, student’s t-test, or two-way ANOVA for appropriate studies.

Additional materials and methods are provided in the supplemental document.

## RESULTS

### Acute liver injury induces rapid and negative responses in skeletal muscle

To understand how liver injury affects skeletal muscle, we evaluated the acute muscle response in two different models of liver injury. First, a widely used chemical hepatotoxic model involving intraperitoneal injection(s) of carbon tetrachloride (CCl_4_) [27] and second, a surgically induced cholestatic liver disease bile duct ligation (BDL) model [27, 28]. Both models consistently demonstrate muscle wasting reminiscent of clinical observations of liver disease patients [29].

As early as 6 hours following CCl_4_ administration, the expression of ubiquitin ligase Trim63 & Fbxo32, critical regulators of early muscle turnover [29], was markedly increased and persisted for 3 days (FIGURE 1a). Additionally, expression of three transcription factors (Lif, Myod1, and Pax7) essentially required for myogenesis were rapidly down-regulated, whereas ankyrin repeat domain 2 (Ankrd2), a powerful regulator of myogenesis and stress responses, was upregulated (FIGURE 1a). We observed increases in both total Smad 2 & 3 protein 24 hours post-CCl_4_ injection, and increases in phosphorylated Smad2/3 two days after CCl_4_ exposure (FIGURE 1b-c). Smad proteins are intracellular signal transducing proteins known to be regulated by TGFβ family members to regulate myogenesis [30].

**FIGURE 1.**
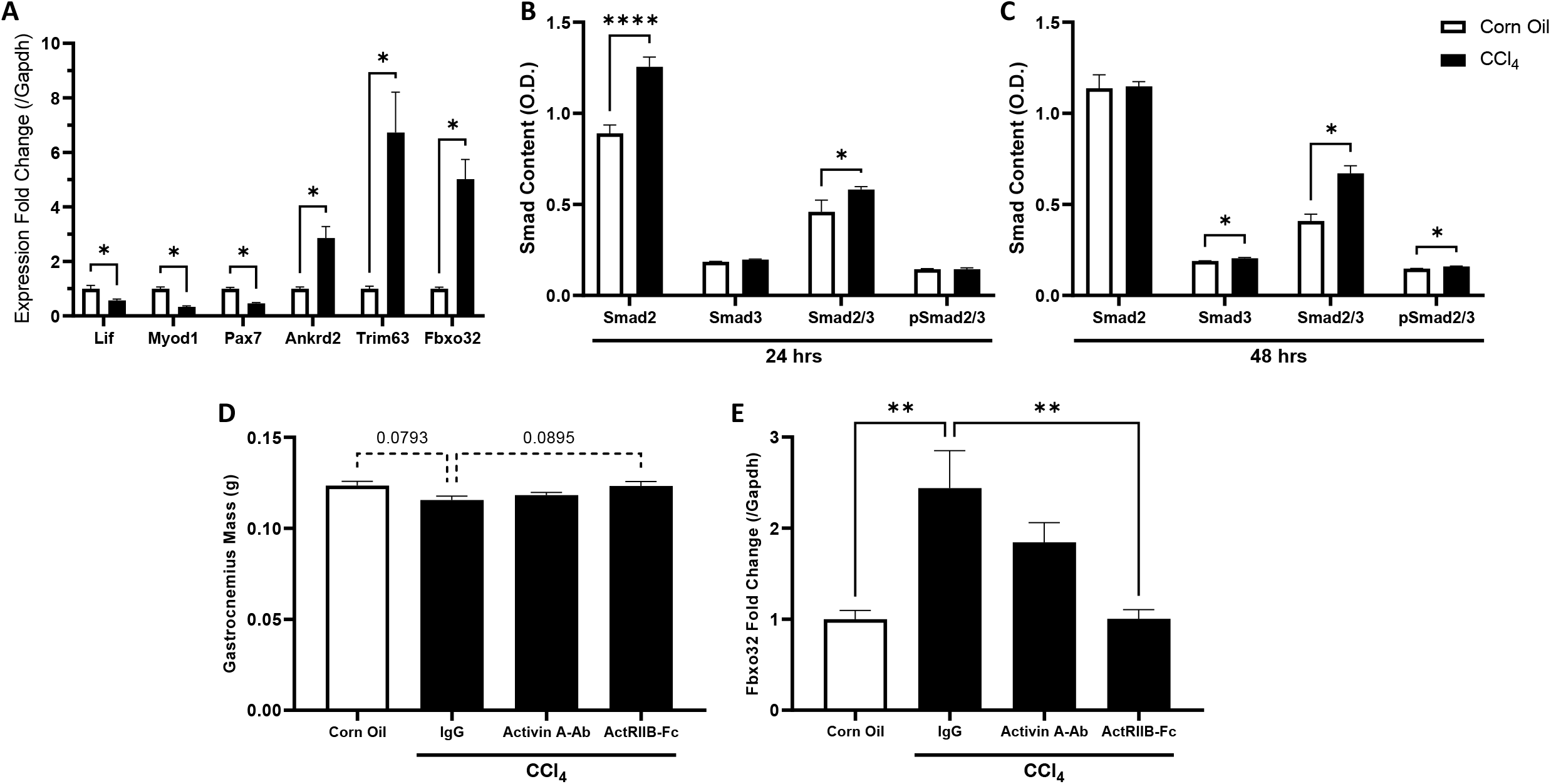
CCl_4_ induced liver injury acutely activates muscle catabolic gene markers and downregulates satellite cell markers. This effect may be mediated through TGFβ signaling, and its effect is ameliorated with ActRIIB inhibition. Female mice were administered a single dose of corn oil or carbon tetrachloride (CCl_4_). Muscle samples were collected at 6, 24, and 48 hours post CCl_4_ injection. **(A)** Muscle mRNA expression 6 hours after livery injury. **(B)** Protein levels of muscle total Smad 2, Smad 3, and phosphorylated Smad 2/3 in muscle as measured via ELISA 24 hours post injury, and **(C)** 48 hours post injury. In a subsequent study, female mice received IgG, Activin A-Ab, or ActRIIB-Fc (10 mg/kg) 16 hours prior to a single administration of CCl_4_. **(D)** Gastrocnemius muscle wet weight and real time PCR analysis of **(E)** muscle Fbxo32 78 hours following CCl_4_ injection are shown. Data are expressed as means ± S.E.M. (n = 5 mice/group). Significance is indicated *P ≤ 0.05, **P ≤ 0.01, or ****P ≤ 0.0001 compared via ordinary two-way ANOVA for figures **A-C** and ordinary one-way ANOVA for figures **D-E.** P-values between 0.05 and 0.1 are indicated by dotted lines

In BDL mice, 24 hours after surgery, ubiquitin ligase (Trim63 and Fbxo32) expression was increased in muscles along with dysregulated myogenic markers (Pax7, Lif, & Myod1), which resulted in lower muscle mass 7 & 14 days after surgery (Supplemental Fig 1a-b).

These data indicate that skeletal muscle sensitively and negatively responds to liver injury through loss of muscle mass and myogenesis suppression in two models of liver disease/injury. These effects have been associated with TGFβ superfamily signaling [22] which were confirmed to be regulated in this context by changes in Smad content. Not only do both models demonstrate muscle atrophy by wet weight and myofiber analysis, but acute gene changes following hepatic insult suggests a mechanism which suppresses satellite cell markers, potentially limiting the muscle’s regenerative capacity. Additionally, the observed increases in Smad2/3 signaling in the CCl_4_ model strongly support a role of a TGFβ superfamily member to be involved in the muscle’s response to acute liver injury. Due to the similar characterizations of BDL and CCl_4_ in skeletal muscle the CCl_4_ model was selectively employed due to its ease of use and reduced surgical complications

### Neutralization of ActRIIB ligands, but not Activin A, prevents the initiation of liver injury-induced muscle atrophy

To ascertain how muscle directly responds to liver injury, primary mouse hepatocytes were damaged via exposure to 0.5% CCl_4_ in culture medium. The concentration was determined as the minimal concentration to cause maximal release of liver enzymes (data not shown). ActRIIB-Fc soluble decoy receptor was added to differentiating C_2_C_12_ myotubes along with diluted injured hepatocyte or control medium with an equivalent CCl_4_ concentration. Culture medium from insulted hepatocytes, but not medium from healthy hepatocytes, inhibited myotube formation (myogenesis) as assessed by reduced myotube diameter. The addition of a ActRIIB-Fc fully rescued myotube formation potential (Supplemental Fig. 2). This observation suggests that injured hepatocytes release ActRIIB-binding ligand(s), including Gdf8, which may exert direct effects on muscle that is not mediated by direct CCl_4_ action on muscle cells. To evaluate liver-muscle communication *in vivo*, female mice were treated with IgG, Activin A antibody (Activin A-Ab), or ActRIIB-Fc sixteen hours prior to a single dose of CCl_4_. Activin A-Ab was selected due to the association of Activin A with CCl_4_-induced liver injury [32]. Three days following CCl_4_ injection, no significant alterations in muscle mass were observed despite statistical trends which may indicate the potential initiation of muscle wasting (FIGURE 1d). Notably, fbxo32 expression was increased with CCl_4_ induced liver injury and blocked by ActRIIB-Fc, but not Activin A-Ab (FIGURE 1e). These data demonstrate that ActRIIB-binding ligands, including Gdf8, potentially modulate the initiation of muscle atrophy after liver injury *in vivo*, while Activin A may contribute little to no biology in this setting.

### Liver injury negatively affects muscle regeneration, which is prohibited by neutralization of ActRIIB ligands

Due to the finding that liver injury causes myogenic satellite cell marker suppression (FIGURE 1a), we queried whether liver injury would adversely affect muscle repair (myogenesis) and, if so, whether TGFβ family members also mediate this effect. Female mice were treated with either IgG or ActRIIB-Fc prior to cardiotoxin (CTX) induced muscle injury with or without CCl_4_-mediated liver injury, which was injected every 3 days for a period of 10 days. A single dose of CTX was injected into the gastrocnemius muscle 6 hours after the first CCl_4_ administration to induce focal muscle injury and muscle repair. As a result, 10 days after CTX injection, simultaneous liver and muscle injury led to skeletal muscle fibrosis and calcification (FIGURE 2a). We observed blunted regeneration in the removal of the necrotic sarcoplasm, followed by concurrent regeneration within the myotube membrane around the necrotic cytoplasm. In addition, prominent collagen deposition and replacement of skeletal muscle tissue with non-muscle cells (FIGURE 2b), and significant reduction in size for nascent fibers (FIGURE 2c-d) were found with liver injury. ActRIIB-Fc treatment prevented these defects in muscle repair. Taken together, we demonstrate that chronic liver injury results in detrimental effects on muscle repair after focal injury supporting our initial findings that suggest a role for hepatic dysfunction to inhibit muscle satellite cell activation and myogenesis. Furthermore, these effects were blunted by ActRIIB-Fc-mediated inhibition of TGFβ members suggesting an extracellular ligand signaling through ActRIIB mediates the liver-muscle crosstalk in this context.

**FIGURE 2.**
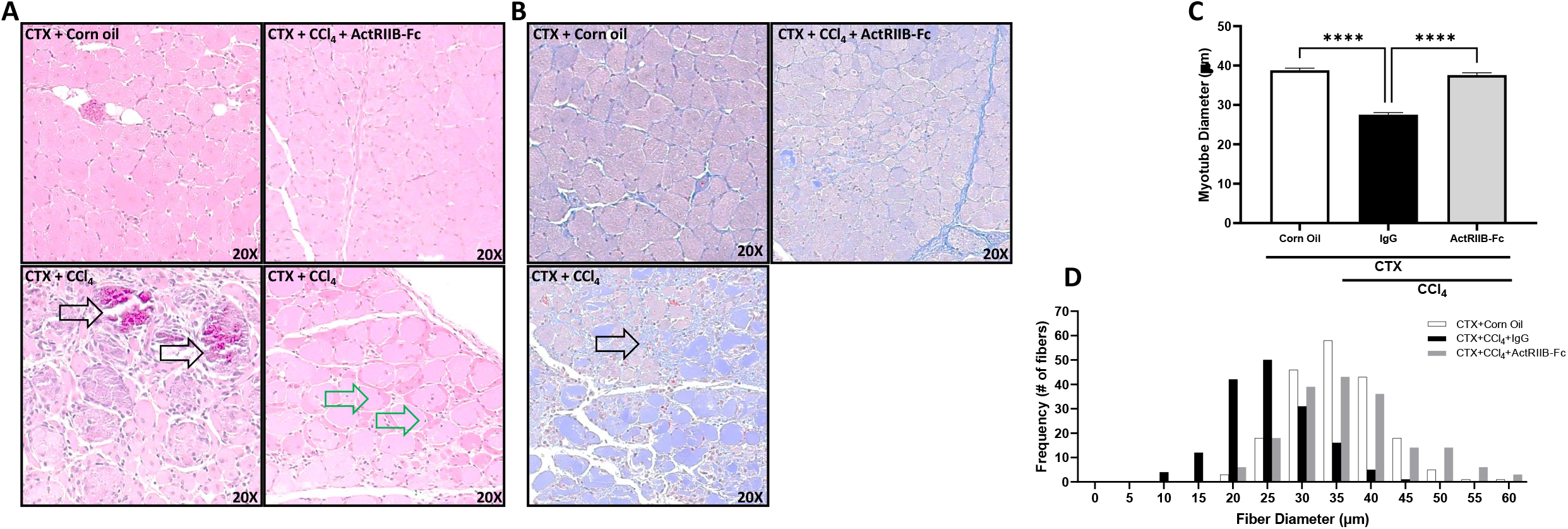
CCl_4_-induced chronic liver injury negatively affects muscle repair following CTX injury. Female mice underwent focal muscle injury via direct cardiotoxin (CTX) injection in their gastrocnemius muscle 6 hours before receiving repeated carbon tetrachloride (CCl_4_) injections over the span of 10 days (3 administrations). An ActRIIB-Fc treatment and IgG isotype control was dosed prior to CTX injections. Muscle samples were harvested at day 10 post-CTX injection. **(A)** Representative H&E cross-sectional images of myofibers in the gastrocnemius muscle. Black arrows indicate areas of calcification (bottom left) and green arrows indicate defective myocyte regeneration (bottom right). **(B)** Trichrome staining of sections (20X). Arrow indicates fibrotic regions. **(C)** Fiber diameter of gastrocnemius muscles and **(D)** the frequency distribution of corresponding fibers (white-corn oil control, black-CCl_4_, and light gray-ActRIIB-Fc). All quantifications of myofibers (~200 counted per group) were determined using ImageScope software (Aperio). Data are expressed as means ± S.E.M. (n = 6 mice/group). Significance is indicated ****P ≤ 0.0001 via ordinary one-way ANOVA

### Neutralization of ActRIIB ligands, but not Activin A, prohibits the progression of chronic liver injury-induced muscle atrophy independent of gender

To elucidate whether ActRIIB ligands mediate progressive muscle mass loss caused by chronic liver injury, male mice were administered CCl_4_ twice weekly for 6 weeks. Twenty-four hours before the first CCl_4_ injection, mice received weekly IgG, ActRIIA-Fc, or ActRIIB-Fc to function as pharmacological ligand traps for their respective receptors. CCl_4_-induced chronic liver injury caused significant muscle mass loss, which was prevented by ActRIIB-Fc, but not ActRIIA-Fc. Treatment with ActRIIB-Fc and ActRIIA-Fc combination did not show additive effects in muscle mass relative to ActRIIB-Fc alone demonstrating an Activin receptor IIB specific ligand is primarily responsible for the induction of muscle atrophy (FIGURE 3a). Interestingly, treatment with both ActRIIA-Fc and ActRIIB-Fc prevented histopathological features of hepatic damage (Supplemental Fig. 3), and ActRIIB-Fc lowered circulating bilirubin levels (FIGURE 3b). ActRIIA ligand inhibition is known to regulate hematopoiesis, which explains the increased bilirubin levels observed in those animals [33].

**FIGURE 3.**
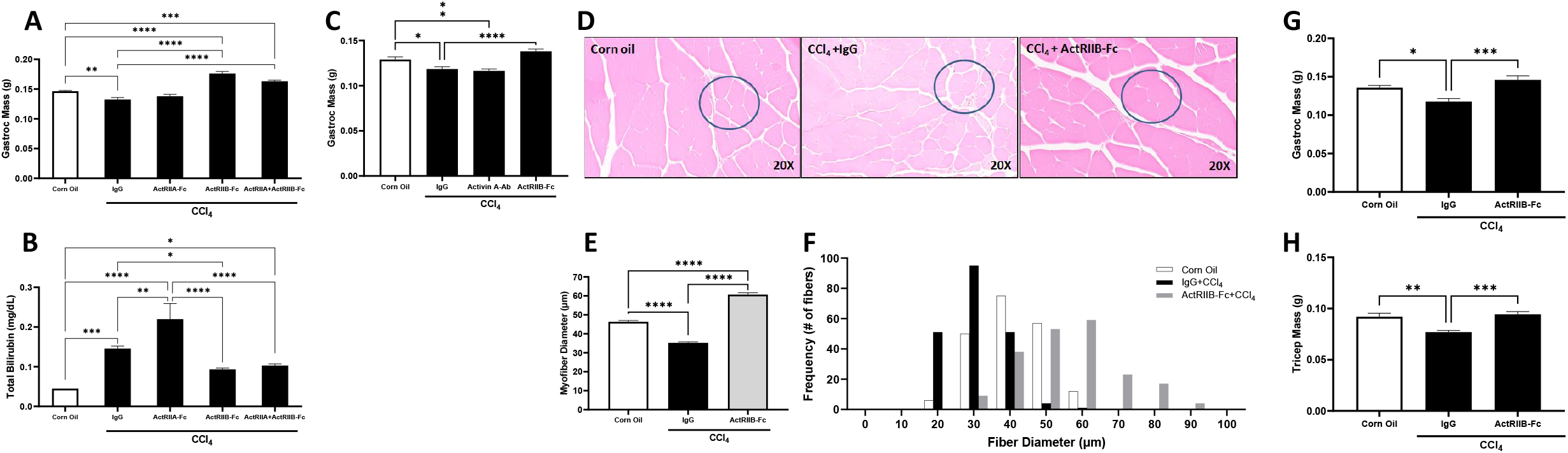
CCl_4_-induced chronic liver injury induces muscle atrophy in mice independent of gender. This effect was prevented and reversed with ActRIIB inhibition which conferred liver functional improvements. Male C57BL/6 mice (n=10) were chronically administered carbon tetrachloride (CCl_4_) twice a week for 6 weeks. Sixteen hours before the 1^st^ CCl_4_ injection each week, male mice received corn oil, immunoglobulin (IgG), ActRIIA-Fc, ActRIIB-Fc, or a combination of ActRIIA-Fc and ActRIIB-Fc (10 mg/kg). A corn oil (non-injured) group was added as a homeostasis control. After 6 weeks of CCl_4_ treatment, **(A)** gastrocnemius wet weights and **(B)** total bilirubin levels were evaluated. In a subsequent study, female mice (n=10) underwent the same CCl_4_ injury paradigm for 6 weeks while receiving IgG, Activin A-Ab, or ActRIIB-Fc weekly. **(C)** Gastrocnemius mass was assessed by wet weight, **(D)** Representative H&E cross-sectional images of myofibers in the gastrocnemius muscle. **(E)** Average fiber diameter and **(F)** the frequency distribution of gastrocnemius muscle fibers. To test the ability for ActRIIB inhibition to prevent muscle degeneration in a pre-existing liver disease state, female mice (n=9) were injected with CCl_4_ or corn oil twice per week for 6 weeks. Then, mice were dosed with ActRIIB-Fc or IgG treatment once per week for 3 weeks during which CCl_4_ or corn oil injections continued (9 weeks total). **(G)** Gastrocnemius and **(H)** triceps brachii muscle masses were assessed by wet weight at study completion. Data are expressed as means ± S.E.M. Significance is indicated *P ≤ 0.05, **P ≤ 0.01, ***P ≤ 0.001, ****P ≤ 0.0001 via Ordinary one-way ANOVA. All quantifications of myofibers (~200 myofibers counted per group) were determined using ImageScope software (Aperio). For figure (F), the Corn Oil bilirubin levels were below the quantifiable detection limits of the assay. Statistical comparison was performed to this group substituting one half of the lower detection limit of the assay

The effects of ActRIIB inhibition to provide protective effects on liver injury-induced muscle wasting independent of gender were confirmed (FIGURE 3c-f) in a 6-week chronic CCl_4_ study in female mice. Based on recent publications, Activin A appeared as the most likely ActRIIB receptor ligand responsible for muscle mass loss in these studies [34], so mice received weekly IgG, Activin A-Ab, or ActRIIB-Fc subcutaneously along with twice weekly CCl_4_ administrations which caused a decrease in muscle mass and myotube diameter which was recovered with ActRIIB-Fc, but not Activin A-Ab. Collectively, our results indicate that (1) chronically-injured liver negatively communicates with skeletal muscle independent of gender; (2) TGFβ family ligands neutralized by ActRIIB-Fc appear to mediate this negative cross talk; and (3) Activin A is not a major mediator in this setting.

### Neutralization of ActRIIB ligands reverses chronic liver injury-induced muscle atrophy

To evaluate the role of ActRIIB ligand inhibition to reverse atrophied muscle after chronic liver injury, ActRIIB-Fc was administered in female mice who had already been injured twice weekly with CCl_4_. After 6 weeks, ActRIIB-Fc or IgG was dosed weekly for 3 additional weeks with continued CCl_4_ injections. Doses and frequency of the molecules were chosen based on previous experiments to elicit optimal increases in muscle mass. Consequently, ActRIIB-Fc-treatment restored muscle mass loss secondary to persistent liver injury (FIGURE 3g-h) compared to controls.

### Neutralization of Gdf8 protects against chronic liver injury-induced muscle mass loss

Another TGFβ superfamily member with a proposed role in liver and muscle communication is Gdf8, which is a myokine expressed in muscle that potently suppresses muscle mass and myogenesis in an autocrine/paracrine manner. Importantly, Gdf8 has high affinity to ActRIIB [35, 36] and this receptor-ligand interaction inhibits proliferation and activation of satellite cells and myoblasts via downregulation of Pax7 and Myod1 expression in muscle [40, 41]. Recent studies have hypothesized a role of Gdf8 in liver disease with concomitant sarcopenia, and is associated with poor survival outcomes in cirrhosis patients [37, 42]. Moreover, Gdf8 was shown to regulate the fibrogenic phenotype of hepatic stellate cells, highlighting a role in hepatic injury response [43].

We assayed human cirrhotic liver samples, and observed significant hepatic Gdf8 protein while undetectable in healthy livers (FIGURE 4a). *In situ* hybridization of fibrotic liver biopsies revealed the source of hepatic Gdf8 to be a subset of hepatocytes, fibrotic cells, and biliary epithelial cells (FIGURE 4b).

**FIGURE 4.**
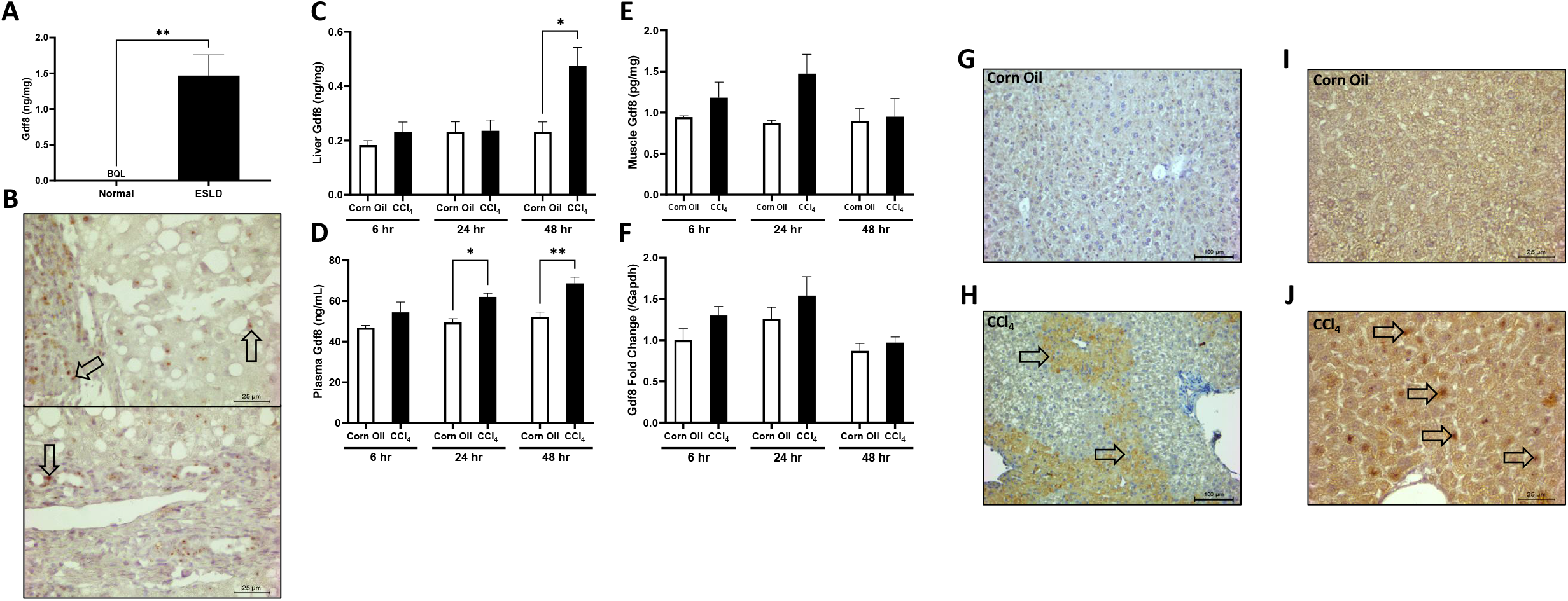
Hepatic Gdf8 content is increased in human patients with end stage liver diseases (ESLD) similar to acutely injured CCl_4_ mouse livers. Liver samples were collected from healthy individuals (n=5) or patients with established cirrhosis (n=7). **(A)** Liver lysates were prepared and subjected to quantification of Gdf8 protein via ELISA. All normal control samples were below quantifiable limits of the assay, and 1 cirrhotic sample was BQL. **(B)** Liver formalin fixed paraffin embedded (FFPE) biopsies from NASH patients were tested for Gdf8 mRNA expression by in situ hybridization. Male mice (n=5) were administered a single dose of corn oil or carbon tetrachloride (CCl_4_).The concentrations of Gdf8 protein was measured via ELISA in the **(C)** plasma, **(D)** liver, and **(E)** muscle at the time points indicated after CCl_4_ injection. **(F)** Muscle gene expression of Gdf8 was measured via RT-qPCR. FFPE liver sections were generated from mice 48 hours post single CCl_4_ injection and stained for Gdf8 via IHC. Representative images of **(G)** corn oil and **(H)** livers are shown. FFPE liver sections were generated from male mice after receiving two injections of CCl_4_ (24 hours apart), and tissues were harvested 24 hours after the second injection. The liver sections were tested for Gdf8 expression via in situ hybridization. Representative images of **(I)** corn oil and **(J)** livers are shown. Data are expressed as means ± S.E.M. Significance is indicated *P ≤ 0.05 and **P ≤ 0.01 via Ordinary one-way ANOVA

Characterization of Gdf8 levels in CCl_4_ injured mice, within 48 hours after a single dose, showed acutely damaged liver produced abundant Gdf8 protein along with increased circulating Gdf8. No significant changes in muscle Gdf8 protein or mRNA were observed (FIGURE 4c-f, Supplemental Fig. 4). Immunohistochemistry of liver sections identified hepatocellular Gdf8 content was increased around central veins (FIGURE 4g-h), while *in situ* hybridization showed induction of Gdf8 mRNA expression in hepatocytes 24 hours following CCl_4_ injury (FIGURE 4i-j). Moreover, we found that the inhibition of C_2_C_12_ myotube diameter by the culture medium from CCl_4_-insulted primary hepatocytes was completely prevented by the addition of anti-Gdf8-Ab to the culture medium (Supplemental Fig. 2).

Hence, Gdf8 appeared to mediate injured liver-muscle crosstalk and thus Gdf8 neutralization may lead to maintenance of muscle mass in the context of chronic liver injury. To test this, female mice were treated weekly with either IgG, Gdf8-Ab, or ActRIIB-Fc, and administered CCl_4_ twice weekly for of 8 weeks. The Gdf8-Ab dose was selected based upon previous experiments identifying the optimal dose in promoting muscle mass. Consequently, we found that Gdf8 neutralization completely protected against CCl_4_-induced skeletal muscle mass loss (FIGURE 5a). ActRIIB-Fc treatment generated more overall lean and gastrocnemius mass than Gdf8 inhibition alone. Interestingly, we found that Gdf8 neutralization reduced hepatic collagen deposition and circulating bilirubin levels to a similar extent as ActRIIB-Fc treatment (FIGURE 5b-c). Furthermore, hepatic Gdf8 protein content was increased with chronic CCl_4_ injury and reduced with Gdf8-Ab and ActRIIB-Fc treatment (FIGURE 5d). To assess the therapeutic potential of Gdf8 neutralization in an existing state of liver disease, we injured male mice twice weekly with CCl_4_ for 5 weeks before initiating concurrent treatment with anti-Gdf8 Ab for 4 more weeks. Compared to IgG controls, anti-Gdf8 therapy significantly recovered the lean mass loss observed in the CCl_4_ injured animals (FIGURE 5f). Gdf8 neutralization reduced hepatic collagen deposition and improved histological markers (hepatocellular rowing, inflammation, biliary hyperplasia) of liver pathology (FIGURE 5g). Collectively, these results demonstrate that (1) injured liver produces Gdf8; (2) and hepatic Gdf8, in part, is responsible for transducing adverse signaling effects of injured liver to skeletal muscle; (3). Finally, Gdf8 also participates in the regulation of hepatic fibrotic response to chronic liver injury.

**FIGURE 5.**
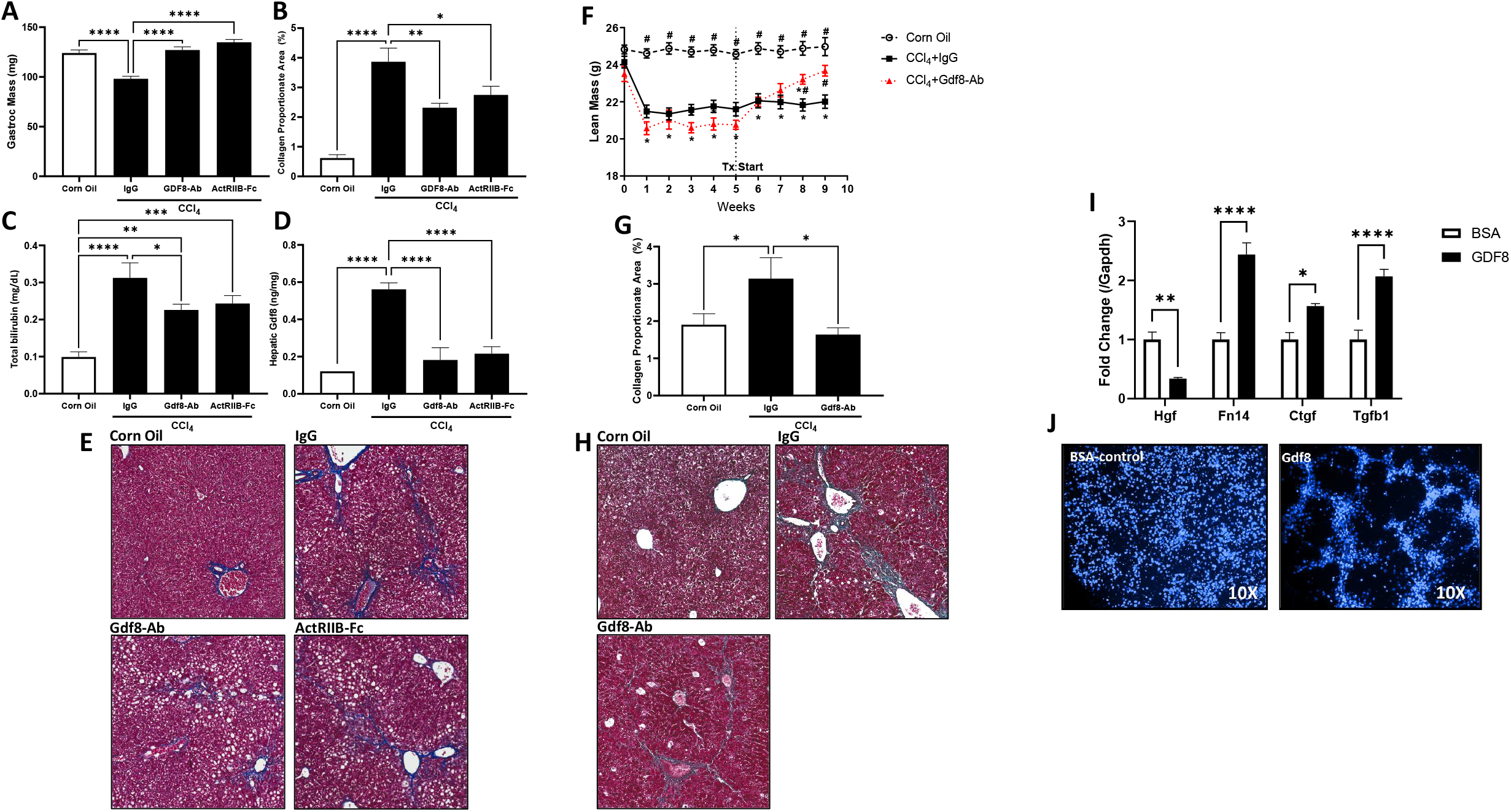
Treatment with Gdf8-Ab or ActRIIB-Fc protects against, and can even reverse, muscle atrophy, reduce hepatic collagen deposition, and lower bilirubin levels in CCl_4_-induced chronic liver injury supporting in vitro data for Gdf8 to function as a fibrogenic factor which activates stellate cells. Male C57BL/6 mice (n=8-10) were chronically administered carbon tetrachloride (CCl_4_) twice a week for 6 weeks. Sixteen hours before each CCl_4_ injection mice received IgG, Gdf8-Ab, or ActRIIB-Fc. A corn oil (non-injured) group was added as a homeostasis control. After 8 weeks of CCl_4_ injury, **(A)** gastrocnemius mass, **(B)** hepatic collagen proportionate area was assessed via image analysis, **(C)** total bilirubin levels were quantified, and **(D)** hepatic Gdf8 levels were measured via ELISA. **(E)**: representative images of Masson’s trichrome staining on liver sections. To test if Gdf8 inhibition can treat existing liver disease, male C57Bl/6 (n=10) mice were chronically administered CCl_4_ or corn oil twice a week for 9 weeks. Starting in week 6, 16 hours before the first weekly CCl_4_ injection, mice received a weekly dose of IgG or Gdf8-Ab with **(F)** continual total body lean mass quantification via QNMR. After 4 weeks of CCl_4_ + antibody treatment **(G)** hepatic collagen content was assessed. **(H)**: representative images of Masson’s trichrome staining on liver sections. To validate fibrogenic role for Gdf8, human hepatic stellate cells (LX-2) were treated with bovine serum albumin (BSA) or Gdf8 protein at equal concentrations (100 ng/mL) for 24 hours (n = 6 wells/group) and **(I)** real time PCR analyses of Hgf, Fn14, Ctgf, and Tgfb1 mRNA expression was performed. **(J)**: representative DAPI stained images (10X) of LX-2 cells treated with bovine serum albumin (BSA) or Gdf8 protein for 24 hours. Data are expressed as means ± S.E.M. Significance is indicated *P ≤ 0.05 compared to corn oil control and # P ≤ 0.05 vs. IgG for LBM QNMR data via repeated measure 2way ANOVA. For all other figures significance is indicated *P ≤ 0.05, **P ≤ 0.01, ***P ≤ 0.001, ****P ≤ 0.0001 via Ordinary one-way ANOVA. Collagen proportionate area was quantified using HALO image analysis software

The response of hepatic stellate cells (LX-2), the major contributor to liver fibrogenesis, to exogenous Gdf8 was also evaluated. Exposure to Gdf8 protein generated a gene expression signature indicative of the stellate cell activation (FIGURE 5i). Additionally, Gdf8 protein stimulated morphological changes in LX-2 cells, redolent of a septa-like structure commonly observed in liver fibrosis (FIGURE 4i-j). These results suggest increased Gdf8 following liver injury may directly act on hepatic stellate cells to promote liver fibrogenesis, supporting reported findings of Gdf8 as a profibrotic agent [43–45].

### Gdf8 disrupts muscle regeneration, mimicking liver injury

Like ActRIIB-Fc, addition of a GDF8 inhibiting antibody to culture media recovered diameter of myotubes exposed to injured hepatocyte media (Supplemental Fig. 2a-b). To test whether liver injury-derived changes in systemic Gdf8 could mediate the negative effect of injured liver on skeletal muscle repair, muscle injury was induced via CTX and, 2 hours later, injected with 5μg of GDF8 protein. The next day, 1μg of GDF8 protein was administered to the injury site. Doses were chosen to mimic the injury-mediated waning in ligand exposure as the protein is quickly absorbed and degraded. Ten days post-CTX injury, muscles treated with Gdf8 displayed a decrease in regenerating myofiber diameter compared to the BSA control group (FIGURE 6), suggesting Gdf8 contributes to liver injury-induced disruption of myofiber regeneration.

**FIGURE 6.**
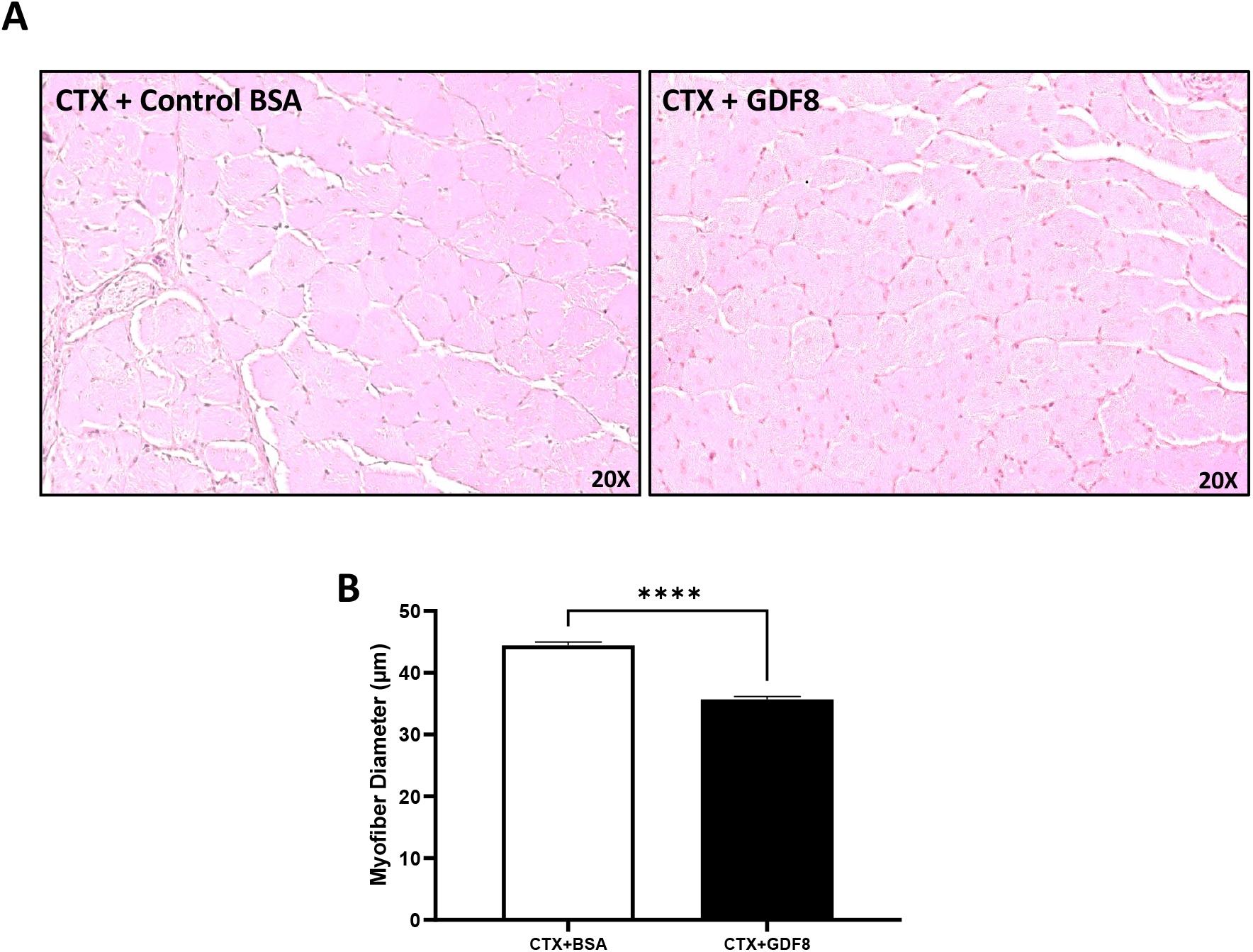
Exogenous GDF8 adversely affects muscle regeneration. Muscle injury was induced by CTX and 2 hours later, 5 μg of Gdf8 or bovine serum albumin (BSA) were directly injected into muscle. The next day 1 μg of each was injected to mimic the injury-mediated waning in ligand exposure. Muscle samples were collected at day 10 post-CTX injury. **(A)** Representative H&E cross-sectional images of myofibers in the gastrocnemius muscle. **(B)** Fiber diameter of gastrocnemius muscles. All quantifications of myofibers (~200 counted per group) were determined using ImageScope software (Aperio). Data are expressed as means ± S.E.M. (n = 5, *P < 0.05). Significance is indicated ****P ≤ 0.0001 via Ordinary one-way ANOVA

## DISCUSSION

These studies demonstrate that ActRIIB-Fc intervention prevents and reverses liver injury-induced muscle atrophy and regenerative capabilities. We show that injured liver produces significant Gdf8 as corroborated by human samples of end stage liver disease. Furthermore, neutralization of Gdf8 largely recapitulates the positive muscle effects of ActRIIB-Fc intervention. Thus, these findings allow us to propose that ActRIIB ligands, primarily Gdf8, are significant mediators transducing the adverse signaling effects of injured liver to muscle. We and others have previously demonstrated that Activin A induces muscle atrophy and degeneration [46]. However, in the context of CCl_4_ induced liver injury, we did not observe overt effects of neutralization of Activin A alone on muscle atrophy. Additionally, in BDL mice, Activin A neutralization did not fully rescue muscle atrophy. These finding suggests that Activin A is not contributing to the injured liver-muscle crosstalk in our preclinical model.

Previous studies by others have suggested several potential mediators connecting liver injury and muscle wasting including hyperammonemia, insufficiency of growth hormone and testosterone [37, 38, 47-51]. Liver dysfunction and portosystemic shunting causes impaired ureagenesis and thus hyperammonemia, a consistent metabolic complication in cirrhotic patients [52, 53]. A NF-kB/myostatin pathway in muscle cells has been proposed to contribute to ammonium acetate-induced muscle degeneration [37]. In addition, growth hormone was shown to inhibit the expression of myostatin in skeletal muscle [54]. These findings propound Gdf8 as a signal mediator downstream of those potential mechanisms. Indeed, cirrhotic patients exhibit increased Gdf8 expression in both skeletal muscle and plasma [55, 56]. Gdf8 is considered as an autocrine/paracrine myokine and a potent suppressor of myogenesis [40, 41]. Here, we demonstrate for the first time that liver injury/fibrosis drives the production of hepatic Gdf8 to negatively regulate skeletal muscle mass. We thereby propose Gdf8 as a candidate hepato-myokine during liver injury. The data prompt the question as to whether Gdf8/ActRIIB ligand signaling is a key central mechanism by which the liver communicates with muscle more broadly in hepatic injury and disease including NASH and cirrhosis.

One intriguing finding from our studies identified a circulatory environment that impedes muscle injury and repair. Our pathological assessment of the combination of liver and muscle injuries unveiled a discovery very redolent of a myopathy where the presence of Ringbinden fibers exists (FIGURE 2A). Ringbinden fibers have been described as an aberrant form of myofibrils that encircle themselves around existing or even dead fibers. Ringbinden fibers have been found predominantly in fast-twitch fibers in mice who possess a mutation in the skeletal muscle α-actin gene (Acta1) [57], which would be consistent in our model where CCl_4_ liver injury selectively affects glycolytic fibers and is reinforced by Gdf8’s ability to affect glycolytic composition and type II fibers [58, 59].

Previously published findings have demonstrated muscle dysfunction in CCl_4_-induced liver injury. Specifically, Weber et al. showed an increase in muscle protein catabolism of rat hindlimb muscles in a manner dependent on severity of liver injury in CCl_4_ injured animals [60]. In that study, the muscle effects of CCl_4_ were limited to fast glycolytic fibers and did not reflect a more diffuse toxic effect in the muscle even in the context of 10-fold increases in CCl_4_ exposure on the muscle due to phenobarbital administration. Additionally, myotoxin induced damage and degeneration can be characterized by specific histopathological features consistent with myofiber damage and immune cell infiltration [61] and this was not observed in our CCl_4_ studies or other publications to our knowledge. These findings from reported by Weber et al. are in line with more recent published data showing increased muscle protein degradation as the mechanism responsible for muscle atrophy observed in a CCl_4_-induced liver injury model [29]. Giusto et al. also reported increases in muscle protein degradation following several weeks of repeated CCl_4_ administration attributable to NF-KB-mediated upregulation of the ubiquitin ligase trim63, which is known to be regulated in muscle by myostatin [62]. Importantly, Giusto et al. also showed reductions in muscle myostatin protein content following several weeks of CCl_4_-induced liver injury. While this may seem initially contradictory to reported clinical reports of increased circulating myostatin levels in liver disease patients who present with muscle atrophy [37, 42], our data suggests muscle may not be the sole source of increased circulating Gdf8 observed in these patients. Ultimately, while we cannot exclude direct effects of CCl_4_ to damage muscle, we do conclude muscle Gdf8 expression is not significantly increased following hepatotoxic injury with CCl_4_ and the observed increased circulating levels must be attributable to primarily liver and potentially other organs.

Giusto et al. also demonstrated differential muscle wasting mechanisms which manifest sarcopenia in two different preclinical mouse liver injury models; bile duct ligation (BDL) and CCl_4_. Therefore, we investigated the potential for a Gdf8 inhibiting therapy to elicit a protective benefit on muscle wasting in a BDL model, which has been shown to drive muscle wasting primarily via Gdf8-dependent mechanisms [62]. Our studies showed a minimal dysregulation of circulating Gdf8 levels or muscle Gdf8 expression following BDL surgery in male ICR mice (FIGURE 7a-b). Despite a lack of statistical significance, Gdf8 protein levels did trend downward at most timepoints, and reached significance 24hrs after surgery, in agreement with previously published findings [62]. Interestingly, despite a lack of substantial increases in circulating Gdf8 following ligation, inhibition of Gdf8 via antibody treatment potently prevented lean mass wasting and myofiber diameter atrophy 14 days post-surgery (FIGURE 7c-d). Like CCl_4_, gene changes in muscles of ligated mice corroborated published findings showing increases in ubiquitin expression and downregulation of myogenic genes Lif and Myod1 (Supplemental Fig. 1). These data suggest not only an increase in muscle protein catabolism following BDL, but a reduced capacity for muscle satellite cells to activate/differentiate to maintain muscle mass in the context of increased catabolism. Importantly, Gdf8 signaling is well characterized to orchestrate these mechanisms [63]. Considering the copious published clinical observations which correlate muscle mass loss to liver disease outcomes, these findings highlight a beneficial effect for myostatin inhibition in liver injury models independent of aberrant hepatic or muscular Gdf8 expression. Indeed, this hypothesis is supported by improved collagen deposition of BDL mice following Gdf8 antibody treatment (FIGURE 7e). A subsequent study evaluated the protective effects of Activin A-Ab and ActRIIB-Fc treatments to preserve muscle mass in BDL mice. Interestingly, Activin A neutralization alone preserved some muscle mass, but not the full protection afforded by ActRIIB-Fc treatment suggesting Gdf8, along with other potential ActRIIB ligands, play a more significant role in muscle atrophy in this model (FIGURE 7f).

**FIGURE 7.**
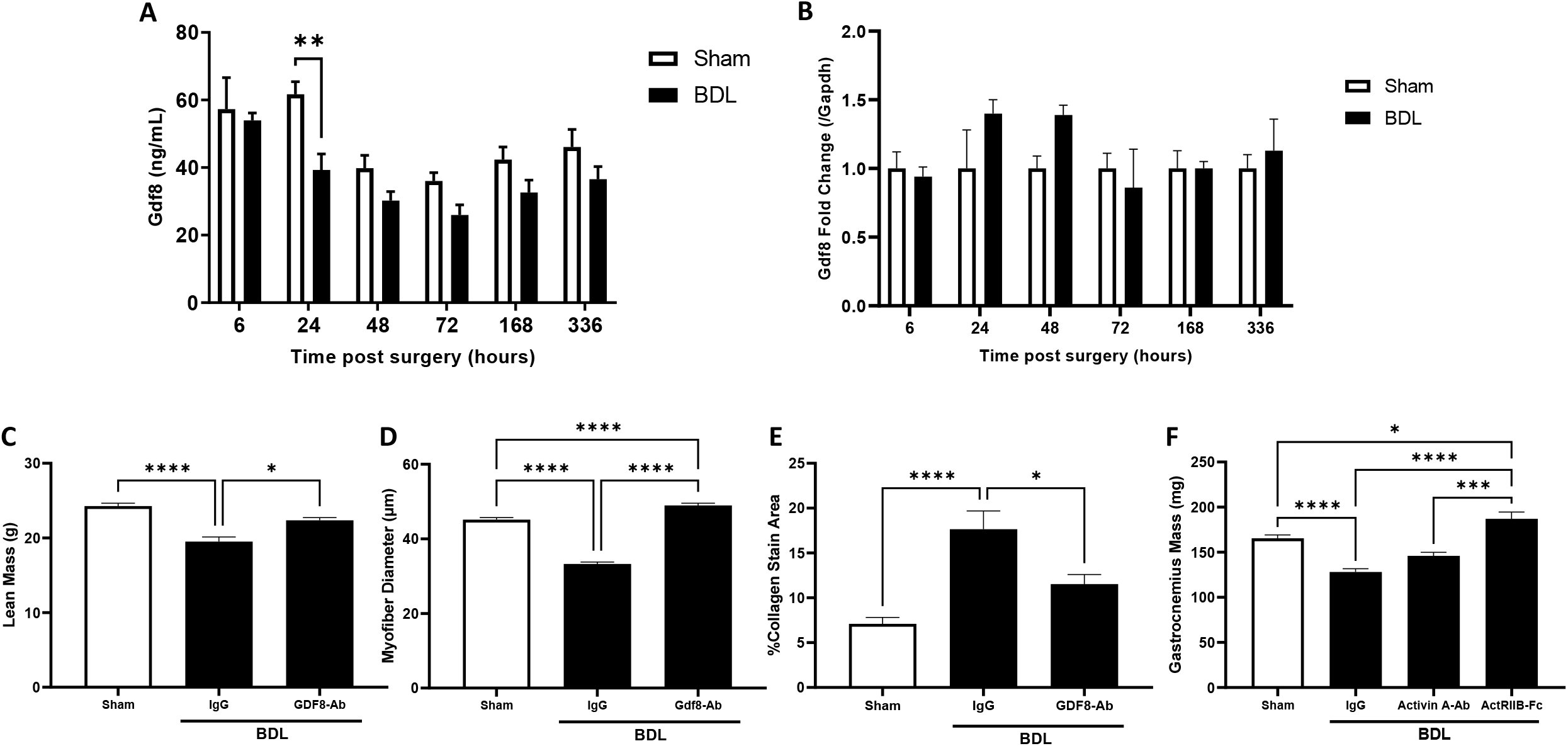
Bile duct ligation model presents with similar lean mass wasting as CCl_4_ which can be prevented with Gdf8 inhibition despite minimal Gdf8 dysregulation which results in lowered hepatic fibrosis. Male ICR mice underwent bile duct ligation (BDL) or sham surgery. Tissues were collected at specified times after surgery. **(A)** Serum Gdf8 and **(B)** muscle Gdf8 expression was characterized. A subsequent experiment involved male ICR mice being pretreated with a Gdf8 inhibiting Ab or IgG before Sham or BDL surgery. **(C)** BDL mice demonstrated potent muscle mass loss 14 days after surgery compared to Sham controls corroborated by reductions in **(D)** myofiber diameter. **(C)** HALO **i**mage quantification of collagen content from BDL and sham liver trichrome stains. Another study pretreated BDL or sham mice with IgG, Activin A-Ab, or ActRIIB-Fc. Fourteen days after surgery, **(F)** gastrocnemius wet weights were reported. All quantifications of myofibers (~200 counted per group) were determined using ImageScope software (Aperio). Data are expressed as means ± S.E.M. (n = 5 mice/group for figure **A** & **B**, n=8 mice/group for figure **C-E**, n=7 for figure **F)**. Significance is indicated *P ≤ 0.05, **P ≤ 0.01, ***P ≤ 0.001, ****P ≤ 0.0001 via 2way ANOVA for figure **A** & **B**, and Ordinary one-way ANOVA for figure **C-F**.

The mechanisms by which Gdf8 contributes to pathogenesis of liver damage are not entirely clear at this time, but multiple studies have shown Gdf8 functions as a profibrotic factor in various tissues [43–45]. Our data has corroborated this role in liver by demonstrating the ability of Gdf8-Ab to reduce CCl_4_ induced liver fibrosis. We postulate that in addition to the endocrine role in inducing muscle atrophy, Gdf8 may play an additional autocrine role by stimulating fibrosis. This was highlighted by the morphological and gene expression changes engendered by Gdf8 that are indicative of stellate cell activation *in vitro* which revealed Gdf8 as a novel and pivotal mediator, which potentially originates from damaged hepatocytes, underlying the potential crosstalk between hepatocytes and HSCs in damaged liver.

Our studies demonstrate inhibition of multiple members of TGFβ superfamily as a promising therapeutic strategy to treat both primary and secondary organ injury or multi-organ injury. Here we demonstrate that by targeting ActRIIB ligands, especially Gdf8, generates dual beneficial effects on both injured liver and wasted muscle. Our findings highlight that in scenarios where liver injury exists, it is vital to not only provide therapies that protect the liver and augment its repair, but also address the resultant muscle atrophy that is a consequence of factors released into the systemic circulation shortly after the liver insult. Here we focused our efforts mainly on evaluating the effects of ActRIIB ligands on muscle degeneration secondary to liver injury. We also revealed that ActRIIB ligands may modulate liver fibrogenesis by directly targeting HSCs. Additionally, our *in vivo* studies demonstrate that Gdf8 inhibition alone does not always compensate for the entirety of muscle effects elicited by pan inhibition of ActRIIB ligands. This leaves open the exciting opportunity to identify additional ligands of therapeutic interest in this disease context.

Based on our findings, we identified a working hypothesis to provide substrate for future investigations to define potential new therapies (FIGURE 8). Recently, cirrhotic patients and the coinciding muscle wasting have been well documented [64–66]. Here we reveal that livers in patients with ESLD also produce abundant Gdf8. Furthermore, we identified specific hepatic cell types in NASH patient liver biopsies that may drive increased Gdf8 expression in fibrotic tissue. It is worth noting that we cannot definitively confirm that increases in Gdf8 observed in those patients is solely attributed to hepatic expression. Indeed, the increases observed in our studies are still relatively low compared to basal Gdf8 expression in muscle, and there are likely additional tissues involved which warrant further investigation. Nonetheless, these findings demonstrate the future potential of therapeutics targeting TGFβ family members for the treatment and control of liver diseases associated with muscle atrophy. Indeed, clinical evidence already exists for this pathway regulating skeletal muscle via ActRIIB-Fc treatment in the form of ACE-031 which has been shown to increase total lean body mass and thigh muscle volume in patients with muscle dystrophy [16]. Furthermore, antibody therapy directed against ActRIIB (BYM338-Bimagrumab) also demonstrates ability to increase lean body mass in the clinical setting. Results from these clinical studies may provide information pertinent to the utility of such therapeutics in liver disease [67].

**FIGURE 8.**
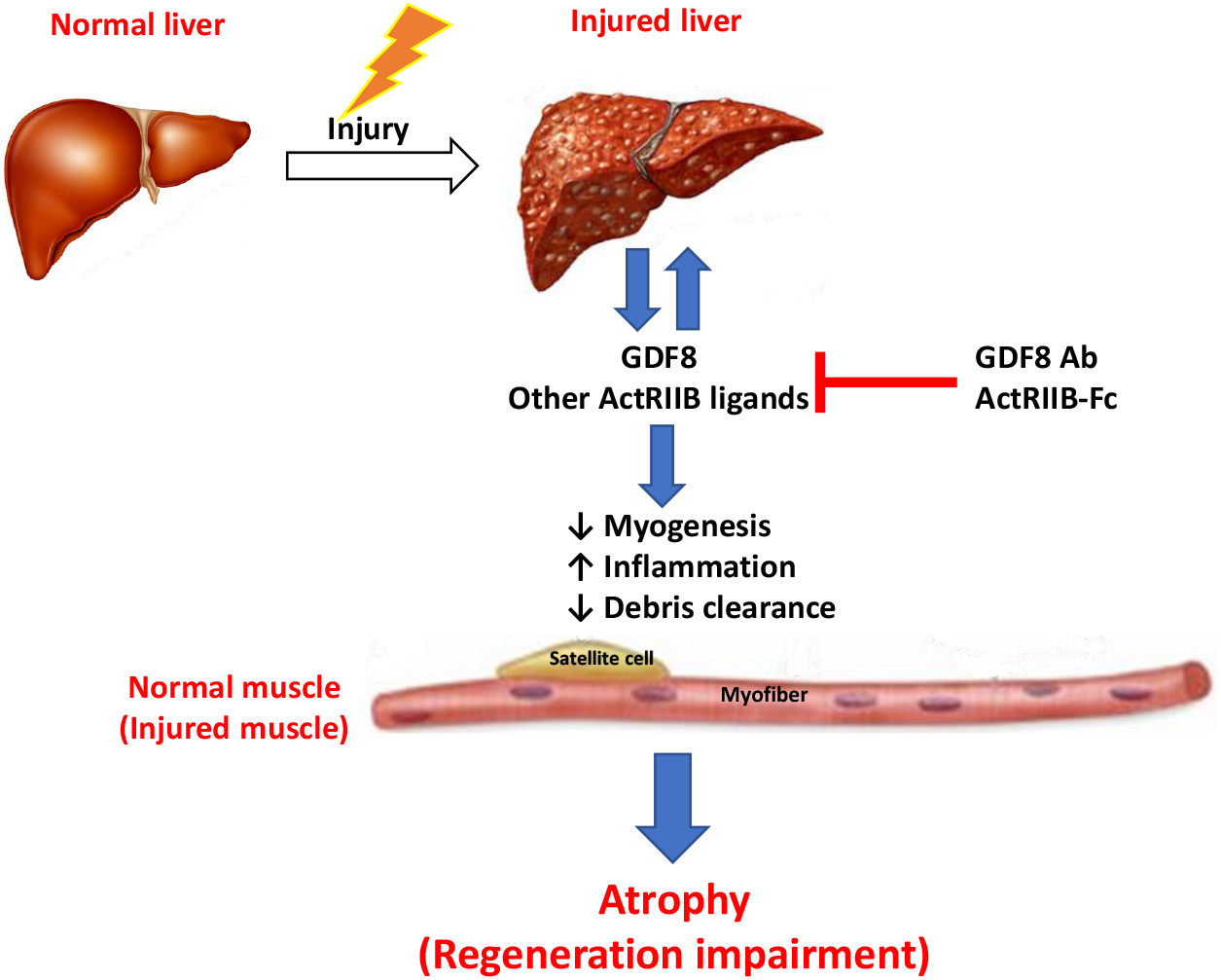
Hypothesis of the liver and skeletal muscle communication under pathological conditions. Injured liver produces and releases Gdf8 and potentially other ActRIIB ligands, promoting liver injury progression and simultaneously causing systemic disruption of TGFβ signaling homeostasis in skeletal muscle and potentially other organs. ActRIIB ligands, mainly Gdf8, induce myofiber atrophy and inhibit myogenesis as well as various biological processes including inflammation and tissue remodeling, resulting in muscle degeneration and regeneration impairment. Degenerating skeletal muscle may in turn negatively feedback onto both the liver and skeletal muscle, promoting the progression of liver injury/fibrosis and concomitant muscle atrophy. Thus, simultaneous inhibition of ActRIIB ligands especially Gdf8 is a powerful approach to prevent or reverse muscle atrophy concomitant to liver injury and even improve both injured liver and degenerating muscle

## ACKNOWLEDGEMENTS

The authors are thankful to Henry Bryant and Ruth Gimeno (Eli Lilly) for the contribution of myostatin antibody. The authors of this manuscript certify that they comply with the ethical guidelines for authorship and publishing in the Journal of Cachexia, Sarcopenia and Muscle [68].

---

**Supplemental Figure 1A.**
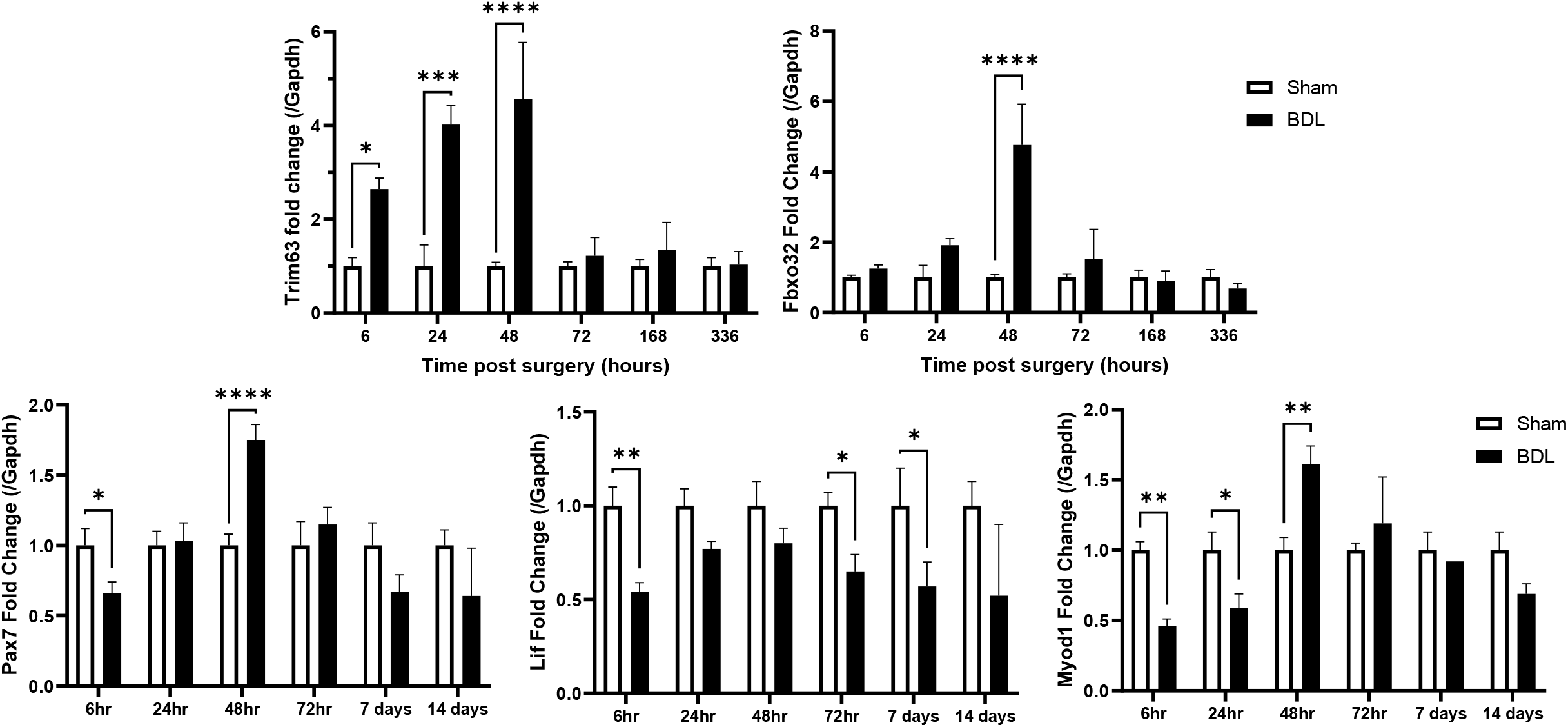

**Supplemental Figure 1B.**
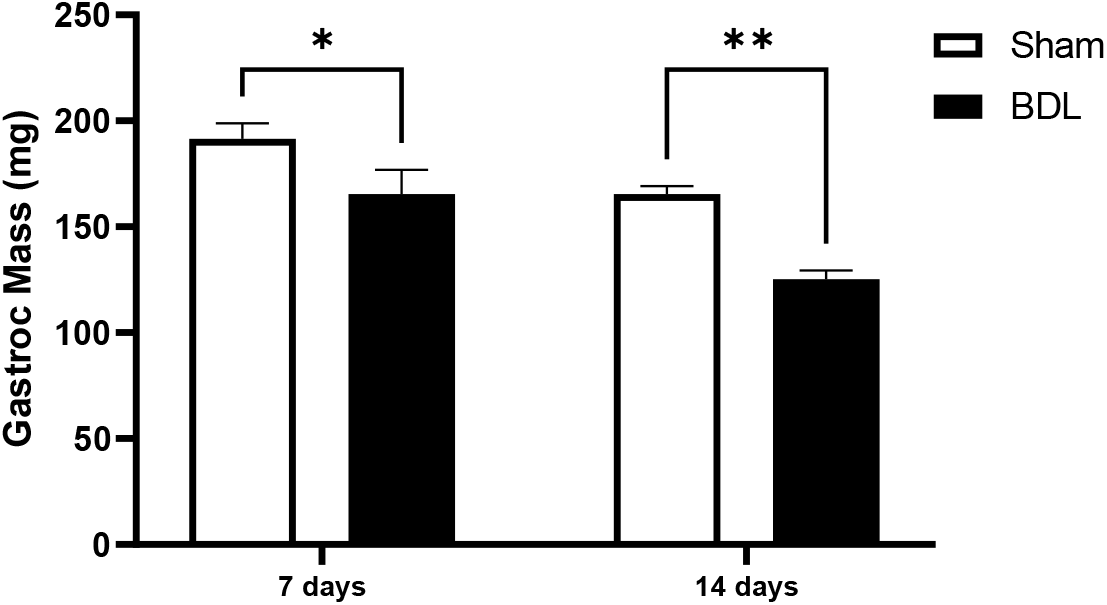

**Supplemental Figure 2.**
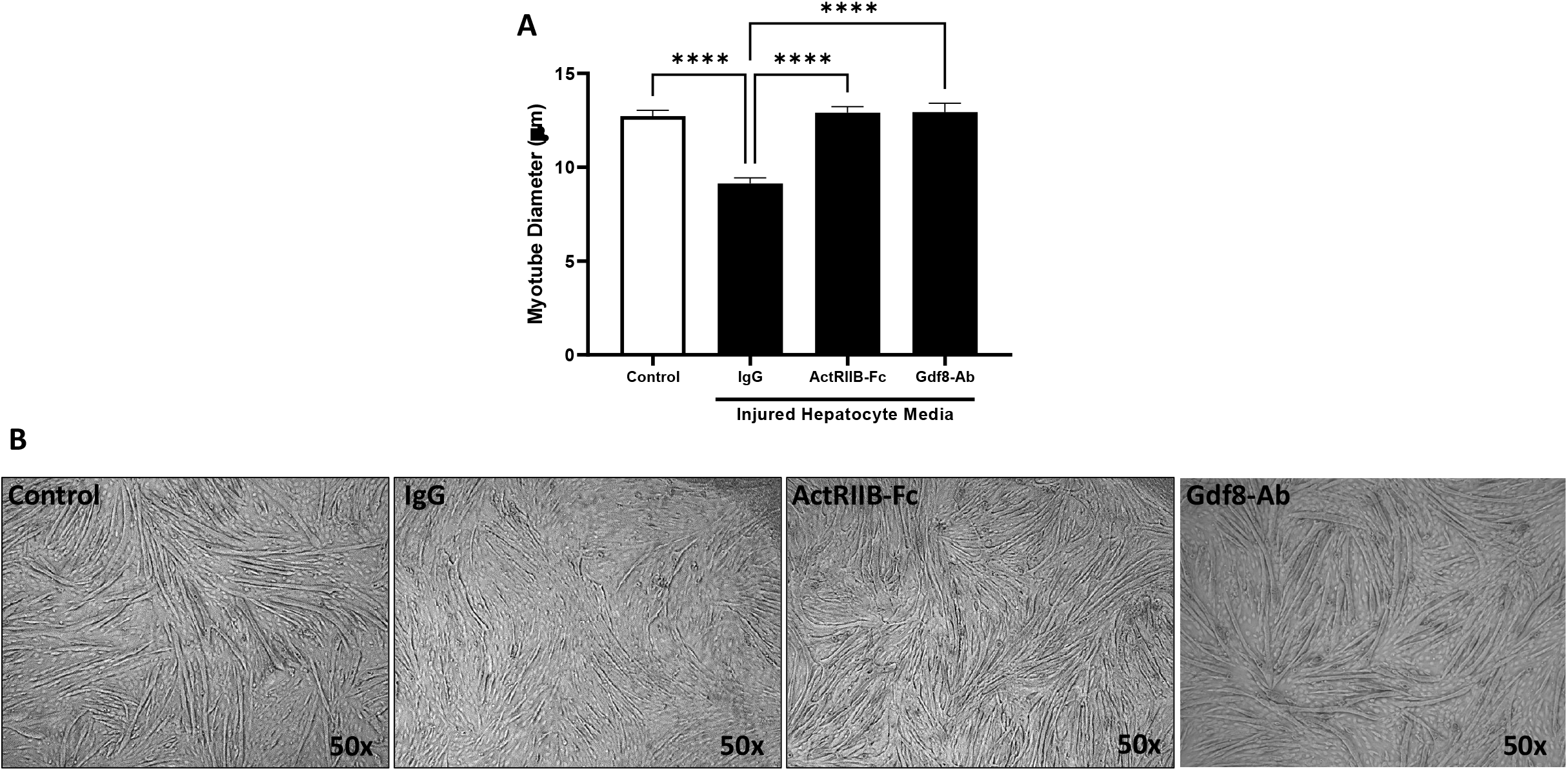

**Supplemental Figure 3A.**
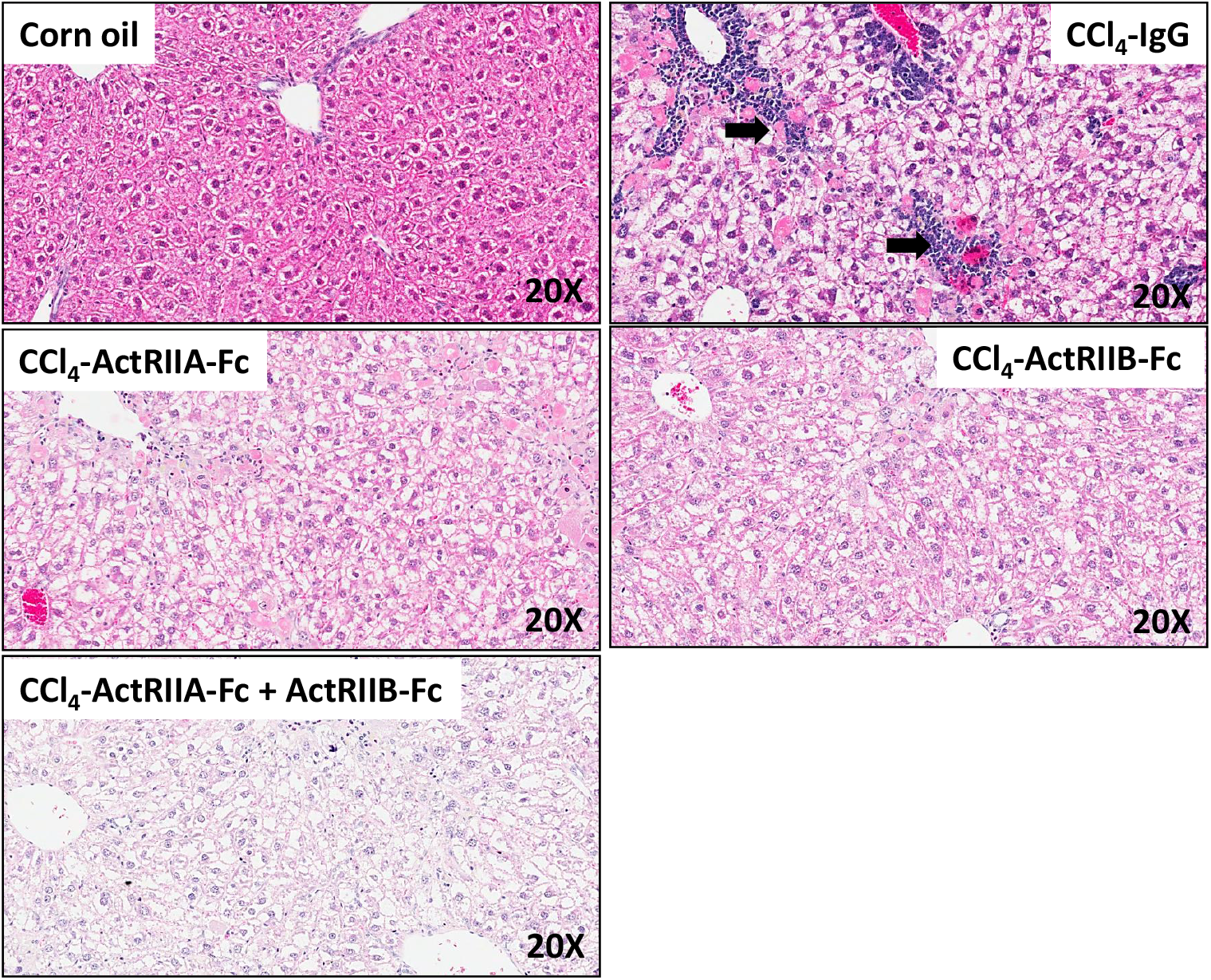

**Supplemental Figure 3B.**
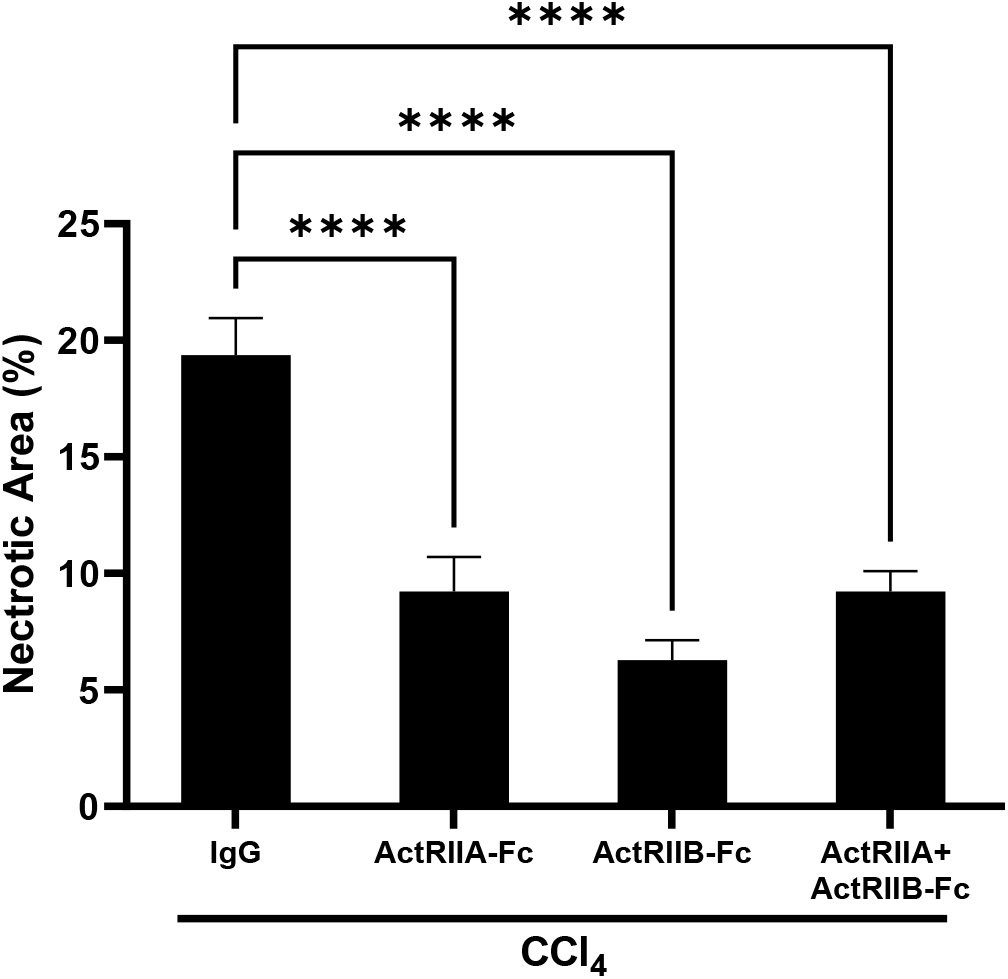

**Supplemental Figure 3C.**
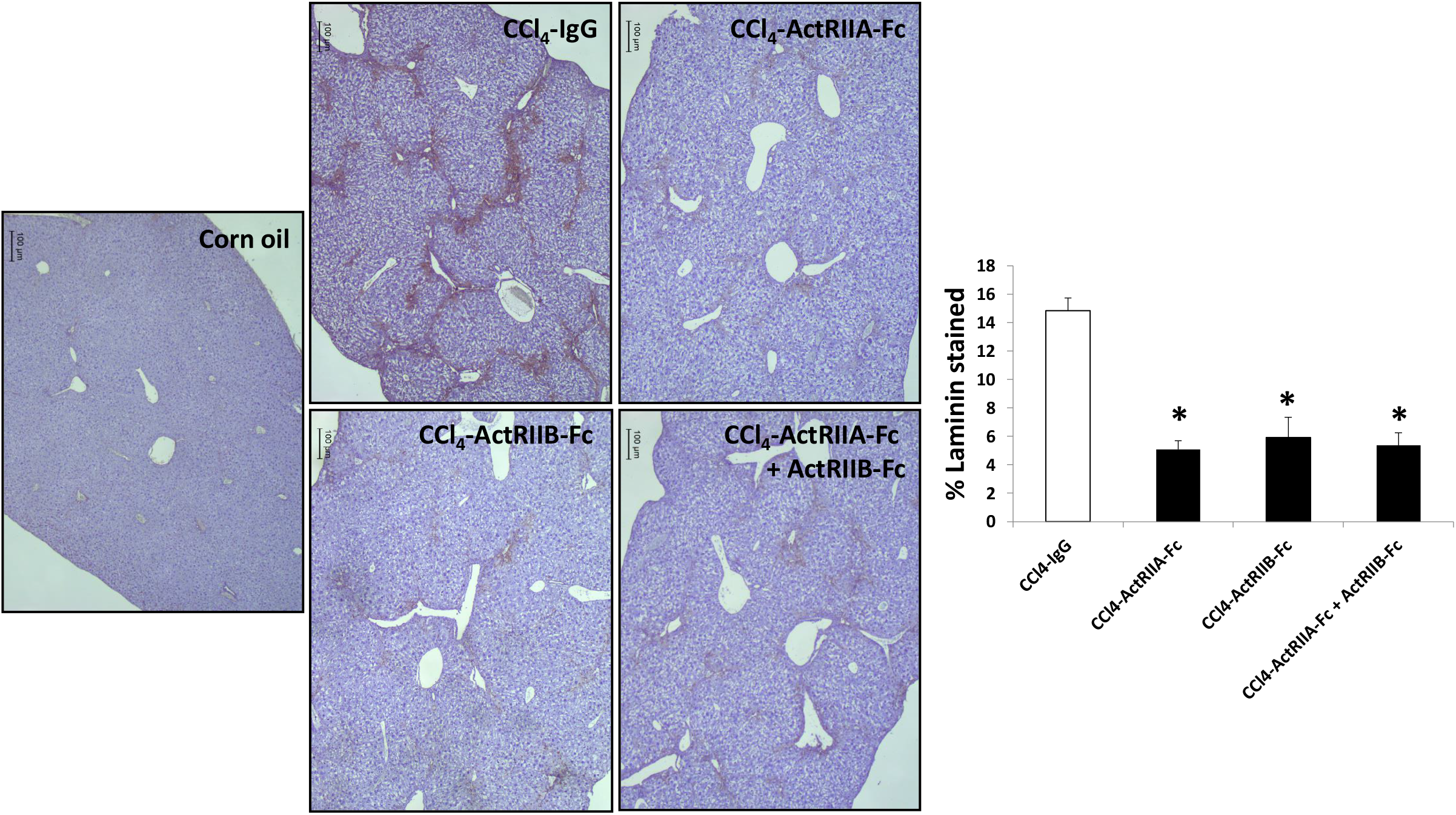

**Supplemental Figure 3D.**
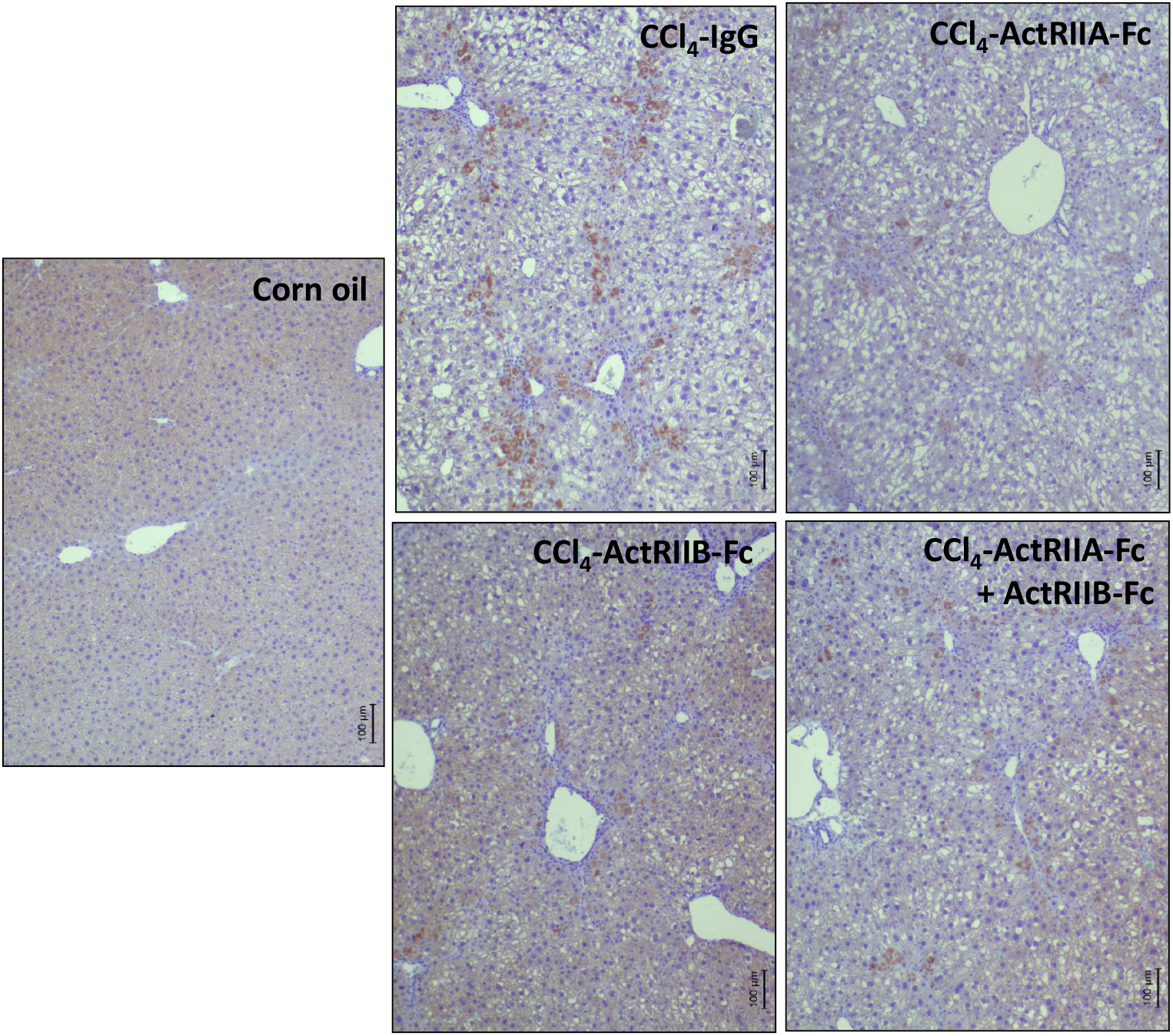

**Supplemental Figure 4.**
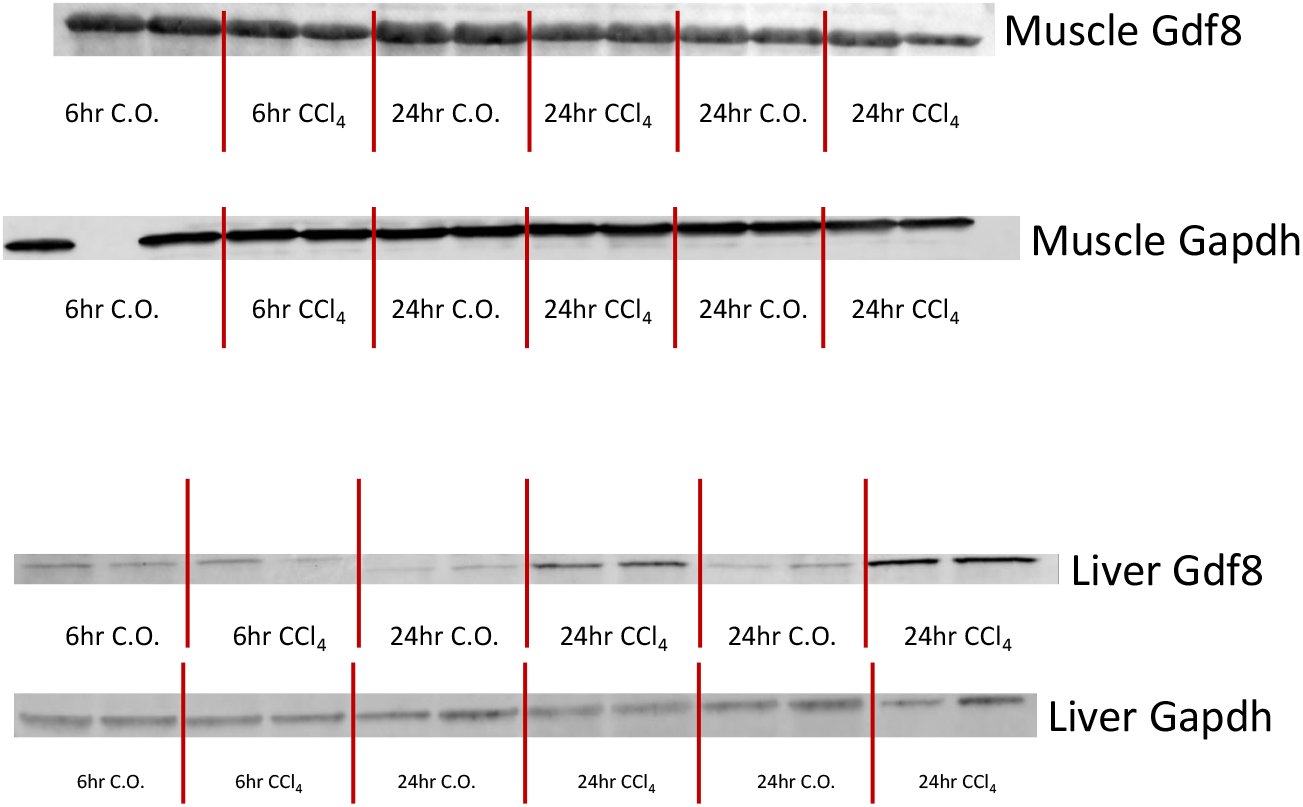

